# A spatio-temporal model for spontaneous thrombus formation in cerebral aneurysms

**DOI:** 10.1101/023226

**Authors:** O. Malaspinas, A. Turjman, D. Ribeiro de Sousa, G. Garcia-Cardena, M. Raes, P.-T. T. Nguyen, Y. Zhang, G. Courbebaisse, C. Lelubre, K. Zouaoui Boudjeltia, B. Chopard

**Affiliations:** Centre Universitaire d’Informatique, Université de Genève, 7, route de Drize, CH-1227 Switzerland; Laboratoire de Médecine Expérimentale (ULB 222 Unit), Université Libre de Bruxelles, CHU de Charleroi, Belgium; CREATIS INSA-Lyon, France; Center for Excellence in Vascular Biology, Department of Pathology, Brigham and Women’s Hospital and Harvard Medical School, Boston, MA 02115; Department of Materials Science and Engineering, Massachusetts Institute of Technology, Boston MA 02139; Unit of Biochemistry and Cellular Biology (URBC), Namur Research Institute for LIfe Sciences (NARILIS), University of Namur (UNamur), 61 Rue de Bruxelles, B-5000 Namur, Belgium

## Abstract

We propose a new numerical model to describe thrombus formation in cerebral aneurysms. This model combines CFD simulations with a set of bio-mechanical processes identified as being the most important to describe the phenomena at a large space and time scales. The hypotheses of the model are based on in vitro experiments and clinical observations. We document that we can reproduce very well the shape and volume of patient specific thrombus segmented in giant aneurysms.

## Introduction

Thrombosis is an important physiological process by which the fibrinogen transported by blood is transformed into a solid and stable mesh of fibrin filaments. Usually, the role of thrombosis is to form a plug on an injured tissue to prevent blood hemorrhage, and to allow for tissue remodeling. Thrombus formation may also have detrimental effects, for instance in strokes, when a clot obstructs a blood vessel without being dissolved by natural thrombolysis. In the case of cerebral aneurysms (fragile, bubble-like structures that may appear on cerebral arteries and cause death if ruptured), thrombosis is the healing mechanism by which the aneurysm cavity may be occluded, and the parent artery remodeled.

Spontaneous thrombosis is found more frequently in giant aneurysms than in smaller ones [23, 20, 6, 13]. A recent approach to treat aneurysms is to insert, in the damaged parent artery, a specific type of stents, called flow diverters (FD). By covering the aneurysm neck, the expected role of FD is to reduce the flow in the aneurysm sac and to trigger thrombus formation in the cavity. Although this treatment has a good success rate, it still fails to produces the expected results in about 20% of the cases.

The great difficulty to understand the mechanisms leading to thrombosis and predict its occurrence, is the interdependence between the biochemical factors, the aneurysm morphology and the blood flow patterns. Computational Fluid Dynamics (CFD) models have been extensively used to provide detailed analysis of flow structures (pressure, velocity, wall shear stress) in aneurysms (see among many others [21, 3]). These numerical simulations do not include any biological processes. However some studies propose an empirical link between flow pattern and thrombus formation. Based on a study with 3 patients, Rayz et al. [19] reported that regions of thrombus formation correspond to slow flow locations. In a second work, the same authors investigated the effect of the flow residence time (FRT) on thrombus presence. They conclude that aneurysm regions with high FRT and low velocity can be prone to thrombus formation. Note that the authors did not consider any molecular processes to justify their claim. A more bottom-up approach is considered in [17, 5], where the flow model is augmented with the presence of abstract pro-coagulant molecules. Despite its simple implementation of the biological response, this model explains very well why spontaneous thrombosis occurs preferably in large aneurysms.

To go one step further, it is necessary to develop more realistic numerical models where, in addition to the flow dynamics, the relevant biochemical molecules involved in the thrombus formation are taken into account, as well as the interactions between each of them and with the rest of the system, typically the endothelial cells and extracellular matrix.

Fogelson and Neeves [9] published a detailed review on the complex interactions between blood components and the vascular wall involved in thrombus formation. The authors highlighted several major points like: (1) The role of Red Blood Cells (RBCs) in the platelets margination from vessels center to the vessel wall surface. (2) The function of the GPIb-vWF bonds in the platelet adhesion at high shear stress. (3) The effect of the hydrodynamic forces on the extension of vWF and the modulation of its biological activity. (4) The complex network of proteins interactions and the polymerization of the fibrin monomers. (5) The way blood molecules are transported by the flow. (6) The threshold density of Tissue Factor (TF) to trigger coagulation. (7) The fact that when the shear stress decreases, the amount of TF needed to generate the same quantity of fibrin decreases. (8) The role of platelets to limit the clot growth by impeding access to TF and to immobilize coagulation enzymes complexes.

Building a fully detailed numerical model which would include all these components, as well as the full coagulation cascade, and could simulate the thrombus formation over the spatial scale of the aneurysms and over temporal scale corresponding to days or weeks is a formidable task, out of reach of current computer technology. For instance, numerical models only implementing the transport of platelets in the presence of the deformable RBCs is already an enormous challenge [14, 15]. Appropriate multi-scale methods are thus necessary. Statistical physics teaches us that many microscopic details may be irrelevant when one considers a process at large scale. The results obtained in [17, 5] clearly show the importance of the spatio-temporal nature of the process, as also emphasized in Fogelson and Neeves [9]. The processes of thrombus start, growth and stop are the result of the presence of the appropriate molecules, at the right place and right time, as transported by the fluid flow. This might be more important than the exact implementation of the coagulation cascade.

In this paper we propose a model to understand the spontaneous formation of a thrombus in a cerebral aneurysm. In section 2 we identify biochemical components and interactions that must be taken into account in view of the time and spatial scales we are interested in. In particular, in-vitro experiments were performed to determine the coagulant factors produced at the aneurysm wall under the flow regime that is observed to be favorable to initiate thrombosis. Then, in section 3 we incorporate these factors on a blood flow model. The main components we consider are thrombin, anti-thrombin, fibrinogen, and fibrin, their mutual interactions, and the response of the aneurysm wall. Other molecules, like platelets are not modeled explicitly, but their role is included in the interaction rules.

In section 5 a validation on two patient-specific aneurysms is performed. It demonstrates the very good predictions obtained from the numerical simulations and illustrates the sensitivity of the process to the production rate of thrombin. Finally this work is concluded and perspectives are given in section 6.

## 2. In-vitro experiments

In this section we present in-vitro experimental results that contribute to the justification of our thrombosis model. Our central hypothesis is that the response of the endothelial cells to low wall shear stress (WSS) flow conditions results into the production of molecules important in the coagulation process, typically Tissue Factor (TF) and its antagonist, the thrombomodulin.

We first consider the synthetic aneurysm geometry shown in Fig. 1 and we numerically determine the WSS on the aneurysm dome when typical physiological flow conditions are applied at the inlet and outlet of the parent vessel.

**Figure 1:**
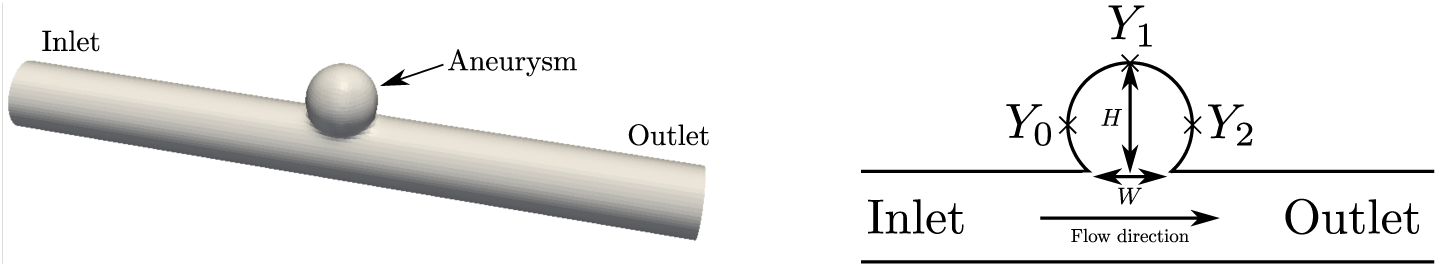
Synthetic aneurysm geometry with the position of the measurement points *Y*_0_, *Y*_1_ and *Y*_2_. The left panel shows a 2D cut of the geometry, in the mid plane along the parent vessel axis.

These WSS conditions are then applied in vitro to a culture of endothelial cells, and the concentration of TF and thrombomodulin is measured. The following subsections describe the above procedure in more details.

### 2.1. Numerical simulation

In order to determine the shear stress to be applied in the in-vitro experiments, a numerical simulation was performed on the simple geometry of Fig. 1. The parent vessel is a straight tube and the aneurysm is a sphere intersecting the tube (the diameter of the parent vessel is the same as that of the sphere and is of 4 mm). The blood is considered as a Newtonian fluid with viscosity *μ* = 4. 10^™3^ Pa s. At the inlet, the time dependent velocity profile depicted in Fig. 2 is imposed, and a zero strain rate in the direction of the flow is imposed as outflow condition.

**Figure 2:**
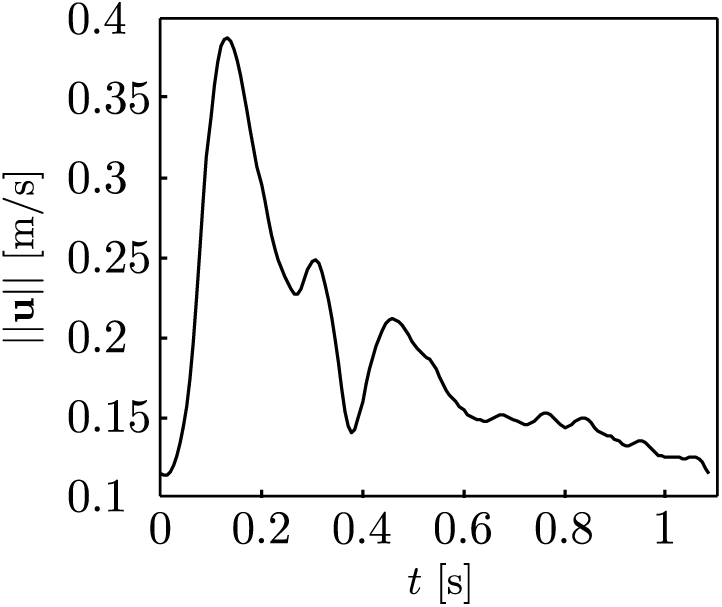
Average velocity profile over one heartbeat imposed at the inlet of the synthetic aneurysm geometry.

The wall shear stress was measured on the three positions *Y*_0_, *Y*_1_ and *Y*_2_, over one period of time, as shown in Fig. 3. These stress signals were then used as inputs for the experimental device described in the following subsections. The simulation was performed with the open source library Palabos (http://www.palabos.org), based on the lattice Boltzmann method which is a validated computational fluid dynamics tool for blood flow (see Závodszky and Páal [24] for example).

**Figure 3:**
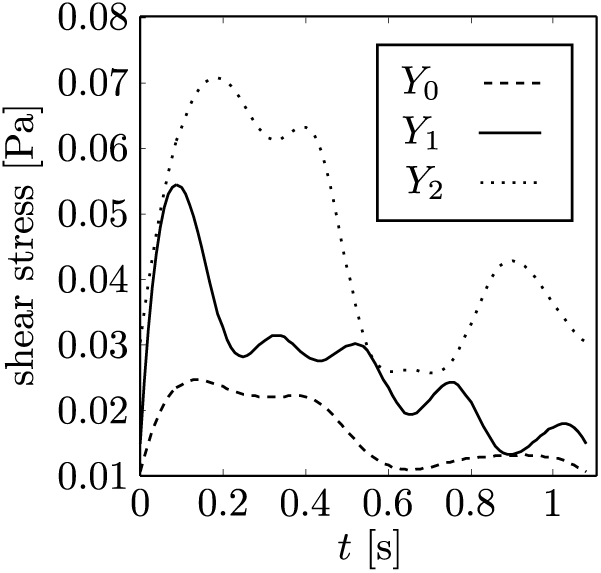
Shear stress over one heartbeat at positions *Y*_0_, *Y*_1_, and *Y*_2_.

### 2.2. Endothelial cells cultures

Human umbilical vein endothelial cells (HUVEC) were isolated from fresh umbilical cords [10] and cultured in Endothelial Growth Medium-2 (EGM-2, Lonza) containing 5% of Fetal Calf Serum. HUVEC were plated at a density of 60, 000 [cells cm^−2^] on gelatin-coated plasma-treated custom plastic plates. HUVEC were then cultured for 24 hours, under different shear stress waveforms (see section. 2.4 for a description of the signals used), or under static (no-flow) conditions. This time duration of 24 hours was selected to let the development of a steady-state regime. RNA was then isolated, and the gene expression are was assessed via polymerase chain reactions (PCR).

### 2.3. Total RNA and Taqman ^®^ low density assay (TLDA) analysis

Following isolation, RNA was reverse transcribed using SuperScript II Reverse Transcriptase (Invitrogen) according to the manufacturer instructions. TLDA analysis was performed with ABi Prism 7000 sequence detection system (Applied Biosystems, Warrington, UK) and data was subjected to relative quantification using 18S (18 S subunit ribosomal protein) as endogenous reference for normalization. The dCt method was applied to each sample in order to calculate the relative fold induction. We focused our attention on Tissue Factor and Thrombomodulin, two important molecules in the coagulation process.

### 2.4. Applying Shear Stress Waveforms to Cultured Human EC

The waveforms prototypical of 3 loci (*Y*_0_, *Y*_1_, *Y*_2_) of intracranial aneurysm cavities (see Fig. 1) were replicated using a custom-made dynamic flow system described in [7, 18, 10]. For the experiments, the flow device was programmed to simulate the waveform corresponding to the particular region of the aneurysmal sac of aspect ratio *AR* = 3 (defined as the height of the aneurysmal sac over the width of the neck, *AR* = *H/W*, see Fig. 1). HUVEC were exposed independently to each of the waveforms for 24h..

### 2.5 Results

The pro-coagulant status of endothelial cells was estimated by the expression (mRNA levels) of Tissue factor (TF), Thrombomodulin (THBD) and the ratio TF/THBD. In Fig. 4 we can observe that the pro-coagulant status significantly increases from *Y*_2_ to *Y*_0_: the ratio TF/THBD is roughly tripled at *Y*_0_ compared to its value at *Y*_1_ and *Y*_0_. This result seems to indicate that the low wall shear stress (WSS) regions on the aneurysm dome are more prone to exhibit a procoagulant behavior. Low WSS is one of the key components of our thrombosis model, as described in the next section.

**Figure 4:**
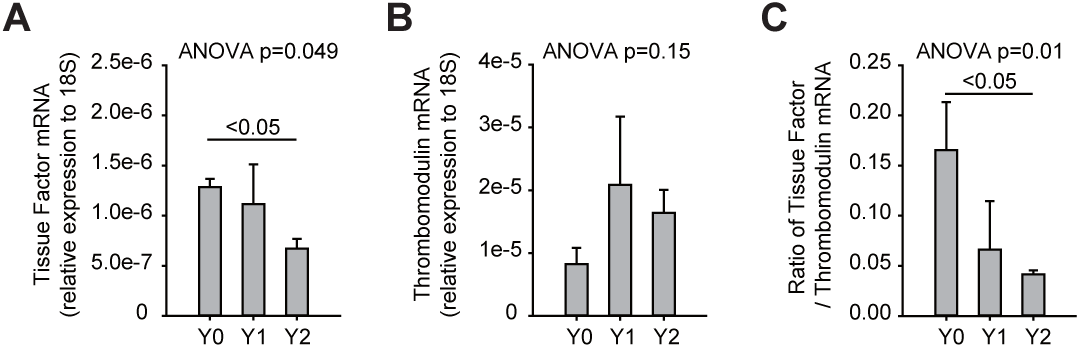
HUVEC mRNA relative production of a) Tissue Factor (pro-coagulant), b) Thrombomodulin (anticoagulant) molecules and c) the ratio between them, detected by PCR, after a 24 hours exposition to corresponding hemodynamic conditions to the ones calculated for *Y*_0_, *Y*_1_ and *Y*_2_, respectively.

## 3 Theoretical thrombosis model

In this section we present the model that we propose to describe thrombosis in cerebral aneurysm. Our goal is to identify a minimal set of components, necessary and sufficient to explain the clinical observations, and in agreement with our current knowledge of the bio-mechanical processes that take place. Simple models are important to improve our understanding by identifying the essential mechanisms. They are also important in order to provide efficient numerical implementations, leading to the possibility of prediction on real cases.

Our model consist in a fluid (the plasma) transporting molecules that can react together, or with the vessel wall, so as to produce or not a thrombus. The molecules considered are: (1) fibrinogen (F), (2) thrombin (T), (3) fibrin (Fi), and (4) anti-thrombin (AT).

Fibrinogen and anti-thrombin are naturally present in blood whereas thrombin and fibrin are created under specific conditions. We assume that thrombin is created by endothelial cells exposed to a low wall shear stress. This assumption reflects the in-vitro observations described in section 2 (the low wall shear stress and residence time hypotheses can also be found in various papers, for example in Chopard et al. [5] and Affeld et al. [1]).

The thrombin released by the endothelial cells can be transported by the fluid, according to an advection-diffusion process. When a thrombin meets an anti-thrombin, both molecules are neutralized. When a thrombin meets a fibrinogen, it produces a fibrin plus a thrombin.

The fibrin is also transported by the fluid but it can stick to another fibrin molecule, or to a vessel wall. By this mechanism, the clot can grow from the aneurysm wall and progress towards the center of the cavity.

Over time, platelets can be attached to the clot and compact the network of fibrin filaments, reducing its porosity. Platelets can possibly cover the entire surface of the clot exposed to the fluid. Once this happen, endothelial cells can regrow [12]. This process explains the layer structure observed in the histology of some clot [16].

A re-endothelialized clot is a stable structure that will reduce the volume of the aneurysm. But if the wall shear stress is still low on this new surface, thrombin will be produced again, and new layers of clot will be formed.

The clotting process stops when the WSS is large enough to prevent further creation of thrombin and when thrombin has been annihilated by the antithrombin. Therefore, the start, growth and stop of the clotting mechanism is deeply related to the flow dynamics, and the interplay between the transport and the reaction processes.

Note that, for the sake of simplification, we do not model explicitly the tissue factor and thrombomodulin. We simply assume that they result in the creation of more or less thrombin. Similarly, the role of platelets is abstracted as a process reducing the porosity of the fibrin mesh and as a factor enabling its re-endothelialization.

## 4 Numerical implementation

### 4.1. General concepts

In order to implement the processes described in section 3, we need to specify many aspects in more details and make several modeling choices. Different implementations are possible but we discuss only one. In our approach, we chose to discretize the simulation domain (parent vessel plus aneurysm) as a 3D regular Cartesian mesh. The time is also discretized in time steps *δt*.

This mesh allows us to implement a flow solver. Here we consider a Lattice Boltzmann (LB) method, coupled with the transport of Lagrangian particles (see below for more details). The blood is assumed to be Newtonian and incompressible. However a Carreau-Yasuda non-Newtonian rheology can be easily implemented.

The Cartesian mesh defines cubical regions of volume *δx*^3^, called spatial cells in what follows. Blood molecules sitting in the same cell at the same time can react according to the processes described in section 3, or can be transported by the plasma across the cells, according to the local flow velocity and a prescribed diffusion coefficient.

The spatial cells give a convenient way to implement the interaction between the blood molecules and the wall, or between the plasma in the wall. The cells of the mesh corresponding to the location of the wall are boundary conditions for the fluid, where a zero velocity is imposed. The growth of the thrombus is modeled by creating new wall cells in the system.

The property of the vessel wall is described as follows. We consider two possible states of a wall cell: the “normal” wall (W) and “thrombosed” wall (TW). We also define the cells close to the walls (CTW) as the fluid (or blood) cells next to a vessel wall (whether W or WT). Flow is possible on a CTW.

The W cells are those covered with endothelium and able to produce thrombin. The TW are the cells next to a W cell which contain fibrin molecules. In a TW cell, no flow is possible and the fibrin forms a solid structure.

Thrombus growth in cerebral aneurysm is assumed to be a very slow process with respect to the time scale of the flow. Thus, the time needed for a cell adjacent to a TW cell to solidify is large. During that time, platelets will have covered the TW cell and enable its re-endothelialization. In other words, a TW cell ends up turning into a W cell, which can produce thrombin and keep the clotting process alive. This is this movement of the wall inside simulation domain that represents the growth of the thrombus.

At each time step *δt* of the model, two main actions are performed successively. First the fluid flow is computed and the particles are transported. Then the reactions are realized, possibly changing the boundary conditions for the fluid.

### 4.2. Particle-particle interactions

The inter-particle coupling correspond to biochemical reactions discussed above. We implement them as a transformation rule, which is applied at each time step of the simulation, with a probability *p*, provided that the corresponding particles are present in the same cell.

The first interaction we consider is the creation of fibrin. It is obtained by the combination of fibrinogen (transported by blood) and thrombin (produced by endothelial cells). This reaction produces fibrin and thrombin with a probability *pF →F i*

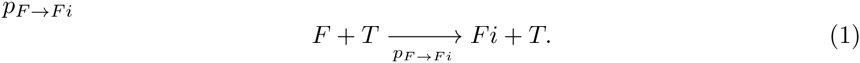

The second reaction of the model is the following: the thrombin and antithrombin annihilate each other with probability 1 when they meet

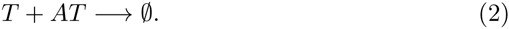

All the other particles do not interact between each others.

### 4.3. Particle-wall interactions

The thrombin is created in CTW cells with probability *p*_*W*_ *→* _*T*_ provided that the adjacent W cells experience a wall shear stress *w* below a critical value *w*_*c*_, for a time larger than a critical residence time *t*_*c*_. These thrombin particles are generated at a random position in the volume *δx*^3^ of CTW cell.

The fibrin can adhere to a blood vessel wall (W or TW) with probability *p*_*Fi→TW*_ provided it is on a CTW cell. This adhesion will transform the CTW cell into a “thrombosed” wall. This TW will in turn be transformed into a W cell, after a characteristic time *t*_*t*_ and with probability *p*_*TW→W*_. The TW cells cannot emit any new thrombin particles before being transformed into a W cell. Finally when a blood cell (B cell) is surrounded by at least four W or WT cells, it can spontaneously clot with a probability *p*_*B→TW*_. Such B cells are called SBW (Surrounded B cell by Wall) in Fig. 6 and in section 4.4.

**Figure 5:**
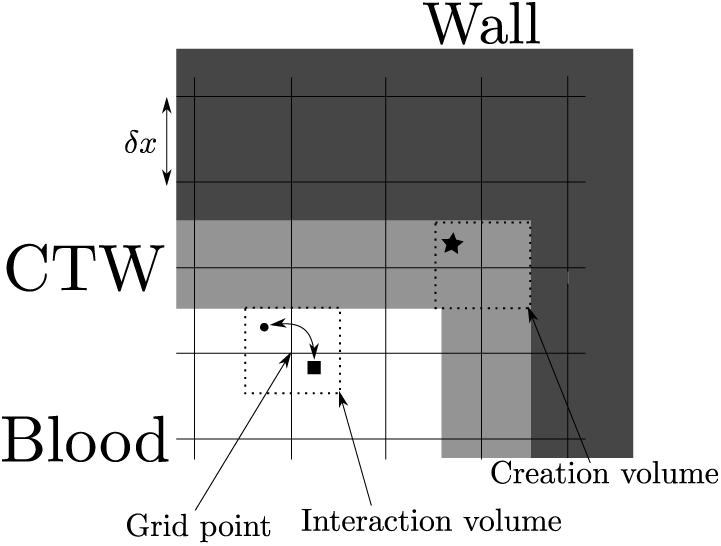
Illustration the interaction range between particles and the different kind of cells. Dark gray cells are the normal wall cells (W), light gray ones are the thrombosed wall cells (TW). The white cells (B) are the cells occupied by blood. The range of interaction is the volume *δx*^3^ around each grid point. The particles represented here as a circle and a square can only interact when located within the same interaction volume. Similarly the “star” particle can only be created in the creation volume.

**Figure 6:**
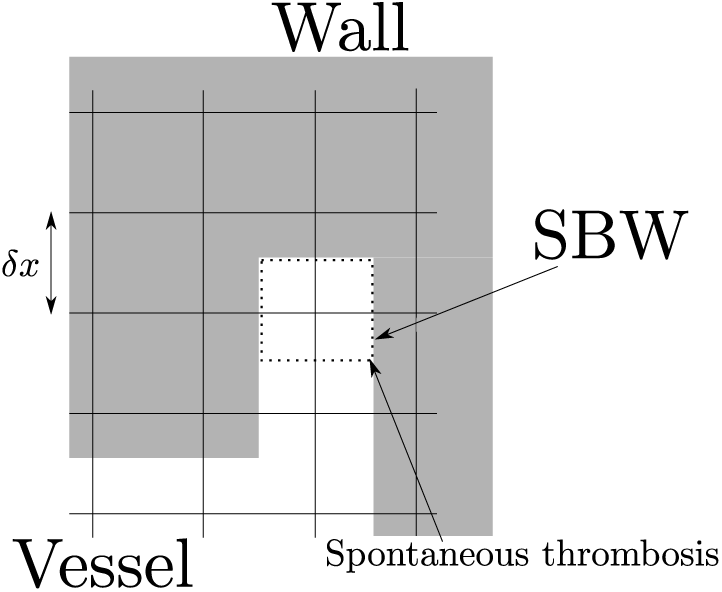
Illustration of the spontaneous thrombosis. When a fluid cell B is surrounded by enough wall cells (thrombosed or normal walls) it can spontaneously be transformed into a thrombosed wall.

### 4.4. Summary of the biomechanical reactions

To summarize, the following creation/destruction/reactions are considered in our model

0. F and AT are naturally streamed by blood.

1. 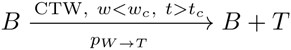
2. 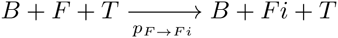
3. 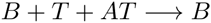
4. 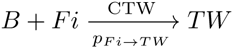
5. 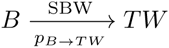
6. 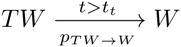

The conditions necessary for a reaction to happen are summarized over the arrows. Each of these reactions occur with certain probabilities (which are indicated under the arrows). For simplicity and in order to accelerate our computations, we took all of them to be equal to one, except for *p*_*W→T*_ which will be varied as shown later.

All these reactions are irreversible and no lysis of the thrombus is included in this version of the model.

### 4.5. Particle-fluid interactions

The fluid is simulated using the lattice Boltzmann method (see Succi [22], Aidun and Clausen [2], Chopard and Droz [4] for more information about the method) and the open source Palabos library. The Palabos library allows the handling of complex geometries and contains a module for particle transport which is used for the simulation of the thrombosis model.

The molecules are viewed as Lagrangian point particles that do not interact directly with the fluid, but are rather modifying the geometry in which the flow occurs. They are transported using a first order Euler scheme. A particle located at a position ***x*** at time *t* will be transported at position ***x***(*t* + *δt*) according to

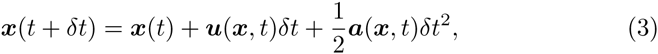

where ***a*** is an acceleration described later, and where

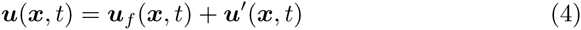

with ***u***_*f*_ (***x***, *t*) the velocity of the fluid at position ***x*** and time *t*. This velocity is obtained by a spatial tri-linear interpolation of the velocity field of the fluid on the neighboring grid points. The velocity ***u****’* is a random velocity which is used to add diffusion into the model. Defining

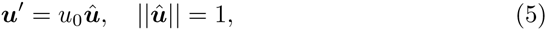

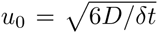 and 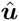 the random direction of the velocity, one can fix *D*, the diffusion coefficient of our particles. Here we assume an isotropic diffusion process. For all the simulations made in this paper the value of the diffusivity is chosen as *D* = 10^−8^ m^2^*/*s in order to create some fluctuations. We did not observe a strong sensitivity to this value.

In order to represent the interaction between the particles and the bloodvessel walls a repulsive force acting on the particles was added. By defining *d* the distance between a particle and the wall, and 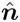 the unitary vector normal to the wall, pointing outwards the computational domain. The force is represented by an acceleration defined as

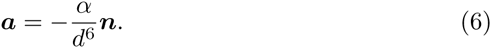

where *α* is a parameter of the model. We chose here *α* = 0.01.

The modeling of the blood molecule could have been performed using a density field instead of explicit point-like particles. The transport would have been obtained by solving a continuous advection-diffusion. Although this approach would have been less expensive in terms of computation power, it was abandoned because of the very small value of the diffusivity constant (*D* = 1^−8^ m^2^*/*s). A diffusivity so close to zero implies the existence of a very thin boundary layer close to the wall surfaces. This boundary layer is therefore very difficult to fully resolve and implies numerical stability issues.

## 5 Validation on two patient specific cases

The validation of the above thrombosis model is a crucial but difficult task. For a proper validation on patient specific cases, many properties of blood, blood-vessels walls, circulation, etc should be known. All these data will have an influence either on the blood flow, on the interparticle reaction rates, on the wall-particles interaction rates, etc. This information is rarely available.

Therefore we will only focus on the ability of our model to reproduce the final thrombus observed in patient specific giant aneurysms. We will first adjust the unknown parameters to obtain the best fit between simulation and clinical observations. Then in order to explore the robustness of the model, some of the parameters will be varied and the effect analyzed.

Note also that segmenting the thrombus from a medical image is a difficult problem. Fortunately, a few cases were produced within the European project THROMBUS (http://www.thrombus-vph.eu).

Our model has been tested on two patient aneurysms which have thrombosed spontaneously (see Fig. 7). For simplicity theses aneurysms will be referred to as *Patient 1* and *Patient 2*, respectively.

**Figure 7:**
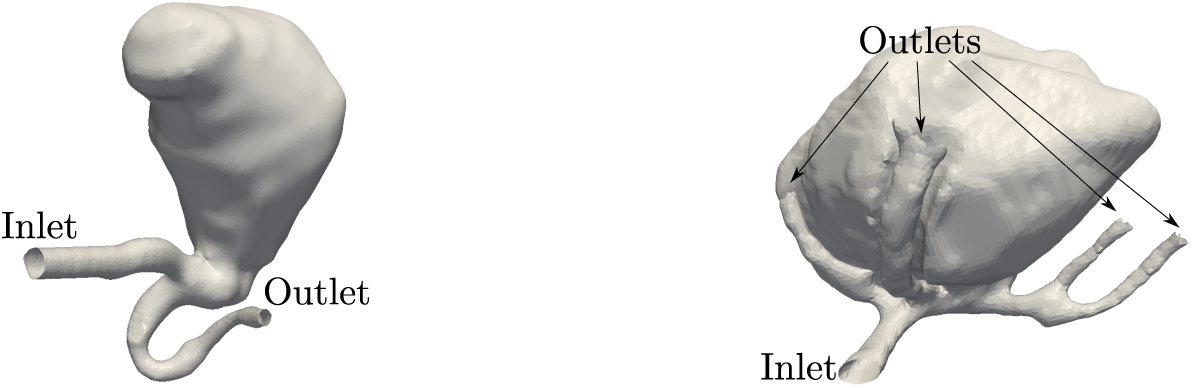
Representation of the giant aneurysms of *Patient 1* (left) and *Patient 2* (right).

For both patients blood flow parameters were based on standard physiological flow conditions, by lack of a better knowledge. The kinematic viscosity is given by *?* = 3.7037 10^−6^ m^2^*/*s, the mean flow rate is *Q* = 4 · 10^−6^ m^3^*/*s, and the density *ρ* = 1080 kg*/*m^3^.

The fluid flow at the inlet is given in Fig. 8 and has a period of one second. The critical wall shear stress is given by *w*_*c*_ = 0.0054Pa, which corresponds to a wall shear rate of 1.35s^−1^, well in the range proposed in [8].

**Figure 8:**
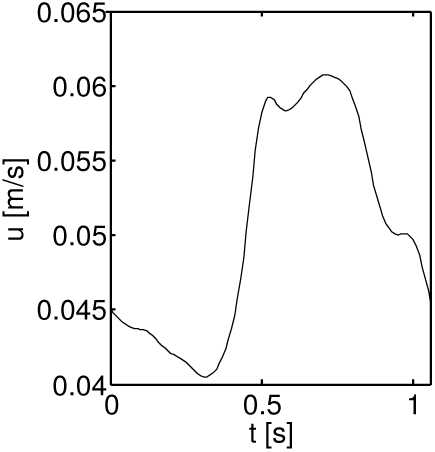
Flow profile imposed at inlet for *Patient 1* and *Patient 2*.

The fibrinogen and anti-thrombin being naturally present in blood, they are injected at the inlets of our simulations, at the rates *Q*_*AT*_ and *Q*_*F*_ given in Table 1. Furthermore the following probabilities, introduced in section. 4.2, are chosen to be 1

**Table 1:** The values of the different parameters used for the *Patient 1* and the *Patient 2* simulations. The space and time steps *δx* and *δt*, the probability of emission of thrombin from the wall cells, *p*_*W→T*_, the critical time *t*_*c*_, and *t*_*t*_ the re-endothelialization time. The injection rate of anti-thrombin and fibrinogen, *Q*_*AT*_, *Q*_*F*_ have units of [mm^−3^s].

| | $\delta x$ [m] | $\delta t$ [s] | $p_{W \rightarrow T}$ | $t_c$ [s] | $t_t$ [s] | $Q_{AT}$ | $Q_F$ |
| --- | --- | --- | --- | --- | --- | --- | --- |
| *Pat. 1* | $8.40 \cdot 10^{-4}$ | $3.87 \cdot 10^{-4}$ | $2.29 \cdot 10^{-4}$ | 1 | 5 | 1200 | 2400 |
| *Pat. 2* | $6.77 \cdot 10^{-4}$ | $2.26 \cdot 10^{-4}$ | $6.99 \cdot 10^{-5}$ | 1 | 5 | 1200 | 2400 |

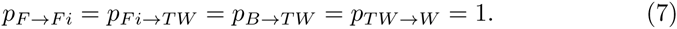

The only probability that is not set to 1 is the emission of thrombin from the blood-vessel walls, *p*_*W→T*_. All the parameters used for the simulations can be found in Table 1. The probabilities *p* used in this paper can also be expressed as reaction/emission/injection rates *Q* by the following formula

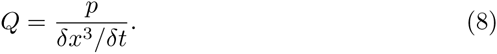

The emission rate of thrombine through the walls of *Patient 1* and *Patient 2* correspond to *Q*_*W→T*_ = 1 mm^−3^s.

The validation is done by comparing the lumen obtained by the simulation with the lumen observed in each patient after natural thrombosis has occurred.

As one can see from Fig. 9 the result for *Patient 1* are in very good agreement. The lumen of the simulation and the results obtained from the clinical data are very similar. We only see a slight underestimation of the size of the thrombus near the neck of the aneurysm. In the case of *Patient 2* the agreement between simulation and clinical observations is less good (see Fig. 10). While the shape of the lumen inside the aneurysm is similar and the clotted volume is equivalent, the position of the lumen is slightly shifted downwards in the simulation with respect to the actual case. At this stage, only a visual comparison is possible as the segmented medical images do not contain enough spatial information to quantify the difference with enough accuracy.

**Figure 9:**
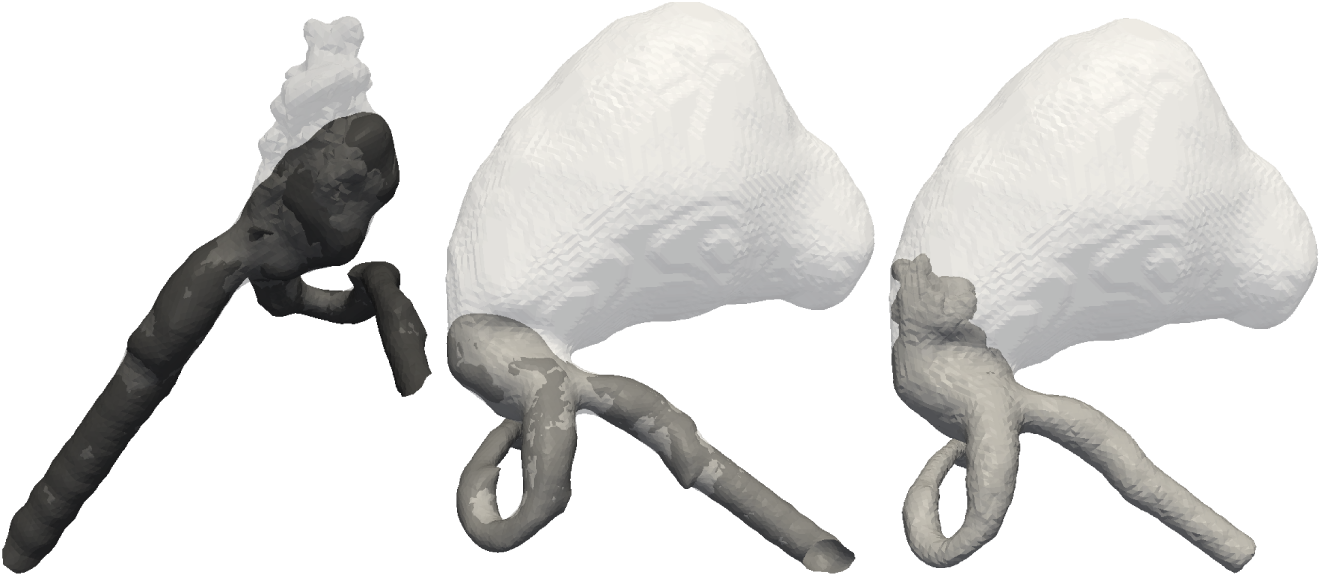
Lumen of *Patient 1* for the real life (middle) and simulation (right) as well as a comparison between the two (left), where in dark gray is the real lumen and in light gray the simulated one.

**Figure 10:**
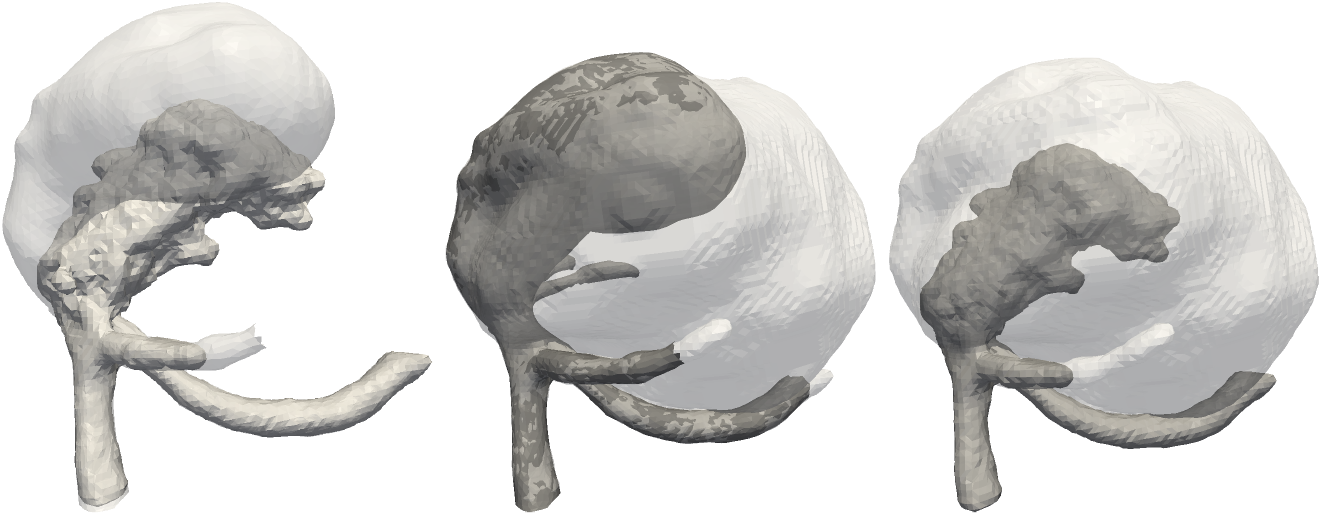
Lumen of *Patient 2* for the real life (middle) and simulation (right) as well as a comparison between them (left). The actual lumen is shown in dark gray and the simulated one in light gray.

The above images correspond to the final stage of the thrombus formation. It is naturally reached in our model as a result of the total annihilation of thrombin by anti-thrombin. As an illustration of the importance of the competition between thrombin production and its interaction with anti-thrombin, we simulated the thrombus formation for three different thrombin emission rates at the walls, respectively 0.01, 0.1, and 1 mm^™3^s.

The results for the two patient geometries are depicted in Fig. 11 and 12. For the emission rates 0.1 and 1, the lumen shape does not change much. As expected, the thrombus progressed only a little bit further for the case of an emission rate of 1 (the light gray region is larger). In the case of an emission rate of 0.01, the effect is more dramatic. Only half of the aneurysm cavity is occluded. The thrombus is stopped much earlier because of the fast depletion of thrombin by the anti-thrombin brought by fresh blood. For *Patient 2* the effect is roughly the same. The size of the thrombus decreases with the increase of thrombin production. However the important change in the volume already occurs between the emission rates 0.01 and 0.1. At an emission rate of 0.01 there is almost no thrombus formation.

**Figure 11:**
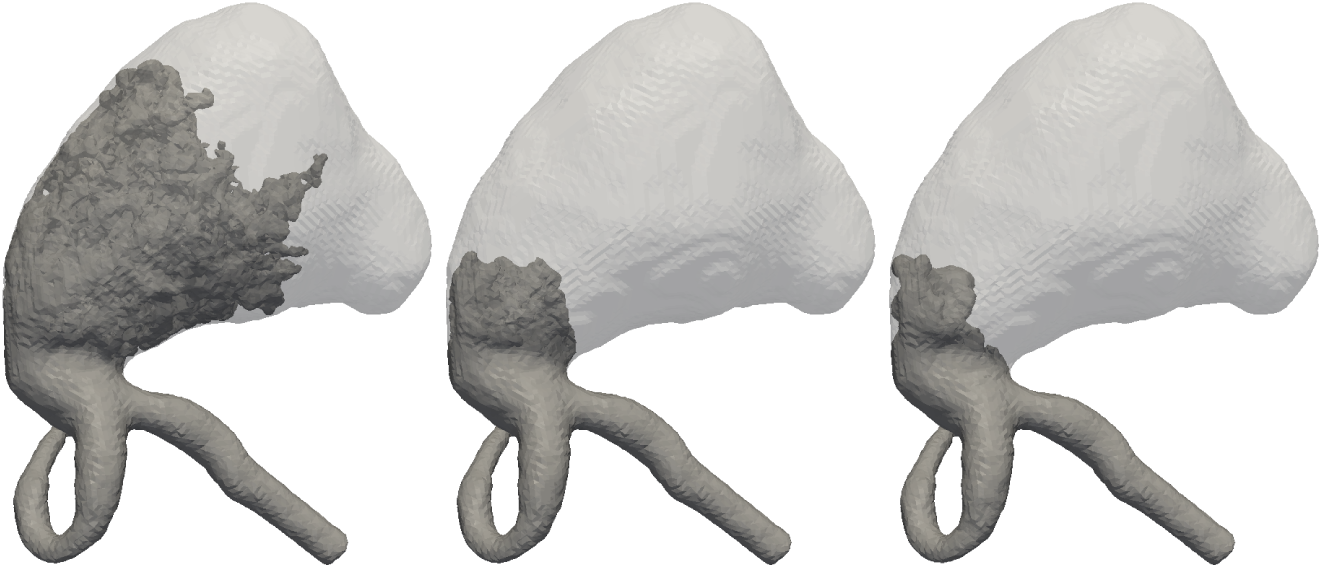
Different lumen shapes (dark gray) for thrombine injection rates of 0.01 (left), 0.1 (middle), and 1 (right) for *Patient 1*. In light gray is the geometry of the aneurysm before the thrombosis.

**Figure 12:**
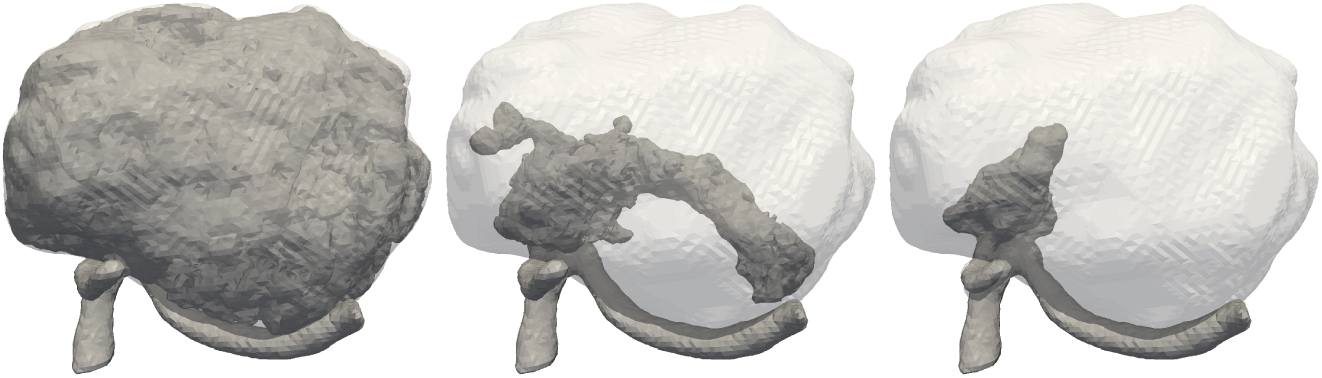
Different lumen shapes (dark gray) for thrombine injection rates of 0.01 (left), 0.1 (middle), and 1 (right) for *Patient 2*. In light gray is the geometry of the aneurysm before the thrombosis.

## 6 Conclusion

In this paper we presented a novel model for thrombosis in cerebral aneurysms, over large spatial and temporal scales. We have proposed a set of biomechanical components that allows us to build a bottom-up numerical approach. The start, growth and stop of the clot in the cavity is an emergent property of the basic interactions present in the system. The main triggering mechanism that we propose is the production of thrombin by the endothelial cells exposed to a low wall shear stress. This hypothesis is confirmed by our in-vitro experiments, and clinical observation [8]. Note that the placement of a flow diverter, whose goal is precisely to cause embolization of the aneurysm cavity by a flow reduction, is also compatible with the hypothesis of a low WSS switch.

The capability of our model to reproduce real thrombi was also corroborated by considering the cases of two patients who underwent a spontaneous thrombosis. We showed that the simulated and actual thrombus have quantitatively the same shape and same volume.

Due to the multi-scale nature of the problem, some strategies have been used to speedup the computation. For instance, amplification [11] is a method by which the reaction rates and the effective size of the particles are increased. By using lower concentrations and by speeding up the reaction processes, we obtained computationally feasible simulations. In particular, the clots shown above were obtained with a few thousands of heart beats, instead of several days or weeks. A drawback of this multi-scale approach is however that the time evolution of the thrombus cannot be captured quantitatively. Only the final stage is significant.

Our results show that the rate of thrombin production at the wall is an important parameters to predict the thrombosed volume. This observation justifies a fortiori the importance of the quantitative measurement proposed in section 2.

Although the results of our simulation are very promising, more work is still needed to better calibrate the model. More patient specific data are also needed, together with in vitro measurements to augment the capability of the model. *Finally, the effect of the drug therapy used during a flow diverter treatment is expected to modify the bio-chemical reaction proposed in our model.* We are currently investigating all these questions.

## Acknowledgments

This work was performed within the European FP7 THROMBUS project. Malaspinas would like to thankfully acknowledge the support of the Swiss National Science Foundation SNF (Award PA00P2 145364). We would like to thank CADMOS for access to supercomputing resources.

## References

[1]. Klaus Affeld, Leonid Goubergrits, Nobuo Watanabe, and Ulrich Kertzscher. Numerical and experimental evaluation of platelet deposition to collagen coated surface at low shear rates. Journal of Biomechanics, 46(2):430–436, 2013. ISSN 0021-9290. doi:http://dx.doi.org/10.1016/j.jbiomech.2012.10.030. URL http://www.sciencedirect.com/science/article/pii/S0021929012006379. Special Issue: Biofluid Mechanics.

[2]. C. K. Aidun and R. J. Clausen. Lattice-Boltzmann Method for Complex Flows. Annual Review of Fluid Mechanics, 42(1):439–472, 2010. doi:10.1146/annurev-fluid-121108-145519. URL www.annualreviews.org/doi/abs/10.1146/annurev-fluid-121108-145519.

[3]. Luca Augsburger. Fluid mechanics of cerebral aneurysms and effects of intracranial stents on cerebral aneurysm flow. PhD thesis, EPFL, Switzerland, 2009.

[4]. B. Chopard and M. Droz. Cellular Automata Modeling of Physical Systems. Cambridge University Press, 1998.

[5]. B. Chopard, R. Ouared, D. A. Ruefenacht, and H. Yilmaz. Lattice boltzmann modeling of thrombosis in giant aneurysms. Int. J. Mod. Phys. C, 18:712–721, 2007.

[6]. José E. Cohen, Eyal Yitshayek, John Moshe Gomori, Savvas Grigoriadis, Guy Raphaeli, Sergei Spektor, and Gustavo Rajz. Spontaneous thrombosis of cerebral aneurysms presenting with ischemic stroke. Journal of the Neurological Sciences, 254(1):95–98, 2007. doi:10.1016/j.jns.2006.12.008.

[7]. GH Dai, MR Kaazempur-Mofrad, S Natarajan, YZ Zhang, S Vaughn, BR Blackman, RD Kamm, G Garcia-Cardena, and MA Gimbrone. Distinct endothelial phenotypes evoked by arterial waveforms derived from atherosclerosis-susceptible and -resistant regions of human vasculature. Proc. Natl. Acad. Sci. U. S. A., 101(41):14871–14876, OCT 12 2004. ISSN 0027-8424. doi:{10.1073/pnas.0406073101}.

[8]. D. Ribeiro de Sousa, C. Vallecilla, K. Chodzynski, R. Corredor, O. Malaspinas, O. F. Eker, R. Ouared, L. Vanhamme, A. Legrand, B. Chopard, G. Courbebaisse, and K. Zouaoui Boudjeltia. Determination of a wall shear rate threshold for thrombus formation in intracranial aneurysms. J. of NeuroInterventional Surgery, 2015. In revision.

[9]. A.L. Fogelson and K.B. Neeves. Fluid mechanics of blood clot formation. Annu. Rev. Fluid. Mech., 47:377–403, 2015.

[10]. G Garcia-Cardena, J Comander, KR Anderson, BR Blackman, and MA Gimbrone. Biomechanical activation of vascular endothelium as a determinant of its functional phenotype. Proc. Natl. Acad. Sci. U. S. A., 98(8):4478–4485, APR 10 2001. ISSN 0027-8424. doi:{10.1073/pnas. 071052598}.

[11]. Alfons G. Hoekstra, Alfonso Caiazzo, Eric Lorenz, Jean-Luc Falcone, and Bastien Chopard. Modelling Complex Systems by Cellular Automata, chapter 3. Springer Verlag, 2010.

[12]. Joseph E. Italiano, Jr., Jennifer L. Richardson, Sunita Patel-Hett, Elisabeth Battinelli, Alexander Zaslavsky, Sarah Short, Sandra Ryeom, Judah Folkman, and Giannoula L. Klement. Angiogenesis is regulated by a novel mechanism: pro- and antiangiogenic proteins are organized into separate platelet alpha granules and differentially released. BLOOD, 111(3):1227–1233, FEB 1 2008. ISSN 0006-4971. doi:{10.1182/blood-2007-09-113837}.

[13]. Z. Kulcsár, A. Ugron, M. Marosfoi, Z. Berentei, G. Paál, and I. Szikora. Hemodynamics of cerebral aneurysm initiation: the role of wall shear stress and spatial wall shear stress gradient. American Journal of neuroradiology, 32(3):587–594, 2011. ISSN 1936-959X. doi:10.3174/ajnr.A2339.

[14]. L. Mountrakis, E. Lorenz, and A. G. Hoekstra. Where do the platelets go? a simulation study of fully resolved blood flow through aneurysmal vessels. Interface Focus, 3(2), April 2013. ISSN 2042-8901. doi:10.1098/rsfs.2012. 0089. URL //dx.doi.org/10.1098/rsfs.2012.0089.

[15]. L. Mountrakis, E. Lorenz, O. Malaspinas, S. Alowayyed, B. Chopard, and A.G. Hoekstra. Parallel performance of an ib-lbm suspension simulation framework. Journal of Computational Science, In press(0):–, 2015. ISSN 1877-7503. doi:http://dx.doi.org/10.1016/j.jocs.2015.04.006. URL http://www.sciencedirect.com/science/article/pii/S1877750315000447.

[16]. Alan T. Nurden. Platelets, inflammation and tissue regeneration. THROMBOSIS AND HAEMOSTASIS, 105(1):S13–S33, JUN 2011. ISSN 0340-6245. doi:{10.1160/THS10-11-0720}.

[17]. Rafik Ouared and Bastien Chopard. Lattice boltzmann simulations of blood flow: Non-newtonian rheolgy and clotting processes. J. Stat. Phys., 121(1-2):209–221, 2005.

[18]. KM Parmar, HB Larman, GH Dai, YH Zhang, ET Wang, SN Moorthy, JR Kratz, ZY Lin, MK Jain, MA Gimbrone, and G Garcia-Cardena. Integration of flow-dependent endothelial phenotypes by Kruppel-like factor 2. J. Clin. Invest., 116(1):49–58, JAN 2006. ISSN 0021-9738. doi:{10.1172/JCI24787}.

[19]. V.L. Rayz, L. Boussel, M.T. Lawton, G. Acevedo-Bolton, L. Ge, W.L. Young, R.T. Higashida, and D. Saloner. Numerical modeling of the flow in intracranial aneurysms: prediction of regions prone to thrombus formation. Annals of biomedical engineering, 36(11):1793–1804, November 2008. ISSN 0090-6964. doi:10.1007/s10439-008-9561-5. URL http://www.ncbi.nlm.nih.gov/pmc/articles/PMC2664710/.

[20]. Bernhard Schaller and Phillipe Lyrer. Focal neurological deficits following spontaneous thrombosis of unruptured giant aneurysms. European Neurology, 47(3):175–182, 2002. ISSN 0014-3022. doi:47978.

[21]. D. Sforza, C.M. Putman, and J.R. Cebral. Hemodynamics of cerebral aneurysms. Annu Rev Fluid Mech, 41:91–107, 2009.

[22]. S. Succi. The lattice Boltzmann equation for fluid dynamics and beyond. Oxford University Press, Oxford, 2001.

[23]. I R Whittle, N W Dorsch, and M Besser. Spontaneous thrombosis in giant intracranial aneurysms. Journal of Neurology, Neurosurgery, and Psychiatry, 45(11):1040–1047, 1982. ISSN 0022-3050. URL http://www.ncbi.nlm.nih.gov/pmc/articles/PMC491643/.

[24]. Gábor Závodszky and György Paál. Validation of a lattice boltzmann method implementation for a 3d transient fluid flow in an intracranial aneurysm geometry. International Journal of Heat and Fluid Flow, 44(0):276–283, 2013. ISSN 0142-727X. doi: http://dx.doi.org/10.1016/j.ijheatfluidflow.2013.06.008. URL http://www.sciencedirect.com/science/article/pii/S0142727X13001379.

